# Adaptive plasticity in plant traits increases time to hydraulic failure under drought in a foundation tree

**DOI:** 10.1101/2020.08.19.258186

**Authors:** A Challis, CJ Blackman, CW Ahrens, BE Medlyn, PD Rymer, DT Tissue

**Affiliations:** Hawkesbury Institute for the Environment, Western Sydney University, Locked Bag 1797, Penrith, NSW 2751, Australia; Université Clermont Auvergne, INRAE, PIAF, F-63000 Clermont–Ferrand, France

**Keywords:** Adaptive capacity, *Eucalyptus*, drought tolerance, genetic adaptation, hydraulic traits, intraspecific variation, phenotypic plasticity

## Abstract

- The viability of forest trees, in response to climate change-associated drought, will depend on their capacity to survive through genetic adaptation and phenotypic plasticity in drought tolerance traits. Genotypes with enhanced plasticity for drought tolerance (adaptive plasticity) will have a greater ability to persist and delay the onset of hydraulic failure.
- *Corymbia calophylla* populations from two contrasting climate-origins (warm-dry and cool-wet) were grown under well-watered and chronic soil water deficit treatments in large containers. Hydraulic and allometric traits were measured and then trees were dried-down to critical levels of drought stress.
- Significant plasticity was detected in the warm-dry population in response to water-deficit, with adjustments in drought tolerance traits that resulted in longer dry-down times from stomatal closure to 88% loss of stem hydraulic conductance (time to hydraulic failure, THF). Plasticity was limited in the cool-wet population, indicating a significant genotype-by-environment interaction in THF.
- Our findings contribute information on intraspecific variation in key drought tolerance traits and THF. It highlights the need to quantify adaptive capacity in populations of forest trees facing climate change-type drought to improve predictions of forest die-back. *Corymbia calophylla* may benefit from assisted gene migration by introducing adaptive warm-dry populations into vulnerable cool-wet population regions.

## Introduction

Rainfall deficits accompanied by warming temperatures are known as “global change-type droughts” and have been observed to cause large-scale tree mortality events around the world (Allen *et al.*, 2010; Dai, 2013). Global change-type droughts are predicted to increase as climate change intensifies (IPCC, 2018), and may drive more tree mortality events in the future. These mortality events impact ecosystem functioning in numerous ways, including alterations to hydrology, and shifts in community structure and composition (Carnicer *et al.*, 2011; Cavin *et al.*, 2013; Vose *et al.*, 2016). Many trees rely on soil water reserves during periods when rainfall is low or absent, but as droughts increase in frequency and severity these soil water reserves may not be replenished (Barbeta *et al.*, 2015). Reductions in soil water content under drought conditions are exacerbated under warm conditions due to an associated increase in vapour pressure deficit (VPD) (Will *et al.*, 2013). Taken together, these changing conditions increase the risk of mortality in trees and may hasten the onset of tree death during drought.

Plants initially respond to increasing soil water deficit by regulating stomatal conductance (*g*_*s*_) to maintain high plant water potentials and slow water loss (Tyree & Sperry 1988; Anderegg *et al.*, 2018). Following stomatal closure (*g*_*s90*_), plants continue to lose water through ‘leaky’ stomata and leaf cuticles (minimum conductance; g_min_; Kerstiens, 1996). Under increasing water stress, the water conducting xylem is placed under increased tension, which can lead to cavitation and the formation of embolisms (air bubbles) that reduce the conductivity of water from roots to leaves (McDowell *et al.*, 2008). If sustained, increasing water deficit may eventually result in total failure of the hydraulic system (Tyree & Sperry 1989; McDowell *et al.*, 2008). Hydraulic failure is widely recognised as a mechanism of mortality in trees under drought conditions (McDowell *et al.*, 2008; Adams *et al.*, 2017; Choat *et al.*, 2018), with critical levels of hydraulic failure in angiosperms commonly being associated with an 88 % loss of stem hydraulic conductivity (expressed as *P*_88_) (Kursar *et al.*, 2009; Urli *et al.*, 2013).

With projected increases in drought-induced tree mortality under climate change, the survival of long-lived plants will depend on their adaptive capacity through genetic adaptation and phenotypic plasticity in drought tolerance traits that delay the onset of mortality (Nicotra *et al.*, 2010; Choat *et al.*, 2018). Studies investigating intraspecific variation in timing of drought-induced tree mortality are urgently needed (Anderegg *et al.*, 2019). Genetic adaptation refers to the trait differences among populations in a common environment. Phenotypic plasticity refers to a change in the expression of a trait measured within or across populations in response to environmental change, while a genotype-by-environment interaction refers to the differential expression of traits by populations to environmental change. Plasticity in cavitation resistance traits at the species level is commonly studied (Holste *et al.*, 2006; Stiller, 2009; Plavcová & Hacke 2012), but less is known about intraspecific variation in plasticity in these traits. Manipulation and common garden studies have provided evidence for intraspecific plasticity in leaf, stem and root cavitation resistance and partitioning of adaptive and plastic variation (Corcuera *et al.*, 2011; Wortemann *et al.*, 2011; Lopez *et al.*, 2013; Claverie *et al.*, 2016; Blackman *et al.*, 2017). One study revealed variable drought tolerance in stem cavitation resistance traits in *Pinus canariensis* populations grown under contrasting water availability in common garden sites with greater drought tolerance observed under drier site conditions (Lopez *et al.*, 2013). Furthermore, a recent review found that plant water availability and variable growth temperatures influence minimum leaf conductance after *g*_*s90*_ (g_min_) (Duursma *et al.*, 2019), and evidence suggests that trees alter their allometry in response to drought, especially with reduced leaf area and increased allocation to roots (Poorter *et al.*, 2012). Taken together, adjustments in these traits in response to water deficit should confer increased drought tolerance and delay the onset of hydraulic failure during severe drought. However, to our knowledge, the level of intraspecific variation in trait plasticity that contributes to short vs long plant desiccation times remains untested.

In this study, we grew saplings of two populations of the south Western Australian tree species, *Corymbia calophylla* (R. Br.) K.D. Hill & L.A.S. Johnson (*Eucalyptus* sensu lato; family Myrtaceae) originating from warm-dry and cool-wet climate-origins under contrasting soil water availability. By examining populations from different climate-origins grown under contrasting soil water availability, we tested for genotype (G), environment (E), and genotype-by-environment (G×E) effects on traits that determine the time it takes for plants to desiccate to hydraulic failure. This study contributes to previous work that suggests potential for adaptive capacity in this species in hydraulic (Blackman *et al.*, 2017), functional (Ahrens *et al.*, 2020), and photosynthetic (Aspinwall *et al.*, 2019) traits. Specifically, Blackman et al. (2017) found greater drought tolerance in a leaf cavitation trait (*P*_50leaf_) in warm climate *C. calophylla* populations relative to cool climate populations and higher *P*_50leaf_ in warm grown saplings in comparison to cool grown saplings. We aimed to measure variability in key hydraulic and allometric traits, including stem *P*_88_ (hereafter referred to as hydraulic failure), minimum rates of water loss (g_min_), total evaporative leaf surface area, and aboveground plant water storage. Saplings were dried-down to estimate the time to hydraulic failure (THF) from the water potential at *g*_*s90*_ (*P*_*gs90*_) to the water potential at *P*_88_ during severe drought. Specifically, we hypothesised that: 1) THF is dependent on a G×E interaction, with longer THF for warm, dry climate populations in response to the water deficit treatment compared to cool, wet populations, 2) THF is genetically determined (G), with longer THF predicted in populations originating from warmer, drier climates compared to cool, wet populations, and 3) THF is environmentally determined (E), with longer THF predicted in trees that show physiological and/or phenotypic adjustment in response to growth under long-term water deficit.

## Materials and methods

### Plant material

The south-west Australian foundation tree species, *C. calophylla* was selected because it is an ecologically important component of forests and woodlands, which have experienced tree mortality and dieback in response to periods of drought and heatwaves (Matusick *et al.*, 2013). Previous studies of this species have shown significant patterns of adaptation to climate (Aspinwall *et al.*, 2017; Blackman *et al.*, 2017; Ahrens *et al.*, 2019a).

Two genetically differentiated *C. calophylla* populations (Ahrens *et al.*, 2019b) with contrasting home-site temperature and rainfall regimes were selected for this study: a high rainfall and cool temperature southern population, Boorara (BOO), and a dry, warm northern population, Hill River (HRI) (Table 1). Seeds were collected from the two populations during 1992 and 2016 by the Western Australian Department of Biodiversity, Conservation and Attractions (previously Western Australian Department of Conservation and Land Management). Seed capsules were sampled from 8 trees located at least 100 m apart and were dried to enable seed extraction. Seeds were stored in a cool room at *ca*. 4 °C. Seeds were germinated (> 80 % viability) and initially grown in forestry tubes in a poly-tunnel at Western Sydney University, Richmond, NSW, Australia.

**Table 1.** *Corymbia calophylla* populations used in this study originating in ‘warm-dry’ and ‘cool-wet’ climate regions with associated mean annual temperature (MAT, °C), mean maximum temperature of the hottest month (MaxT, °C), mean annual precipitation (MAP, mm), mean precipitation of the driest month (PreDM, mm), and 1/aridity index (1/AI).

| Population | Acronym | Climate region | Lat | Long | MAT | MaxT | MAP | PreDM | 1/AI |
| --- | --- | --- | --- | --- | --- | --- | --- | --- | --- |
| Boorara | BOO | Cool-wet | -34.639 | 116.124 | 15.0 | 25.6 | 1159 | 24 | 0.95 |
| Hill River | HRI | Warm-dry | -30.311 | 115.202 | 18.8 | 31.7 | 563 | 4 | 2.56 |

### Experiment set up and design

*Corymbia calophylla* saplings underwent four phases during the experiment. The initial phase was the *establishment phase* in which saplings were grown under well-watered conditions for 2.5 months. The second phase was the *treatment phase* in which saplings were grown under either a well-watered (W) or a water deficit (D) treatment for 4 months. The third phase was the *drought hardening phase* which was necessary to avoid rapid mortality in well-watered saplings in the final *drought to hydraulic failure phase*. The final phase was the *drought to hydraulic failure phase* in which water was withheld from saplings until they reached the water potential inducing 88 % loss of conductivity (*P*_88_). Saplings were then harvested.

Saplings were grown in a poly-tunnel facility situated on Hawkesbury campus, Western Sydney University (Richmond, NSW, Australia). The poly-tunnel facility contained 91 pallets and population × treatment combinations were randomly distributed to minimise microclimate affects. Two rows of buffer saplings surrounded the experimental saplings to minimise edge effects.

### Establishment phase

On 30^th^ August 2017, 64 similar sized saplings from each *C. calophylla* population (64 BOO, 64 HRI, 128 total) were transplanted into 45 L woven plant bags (one sapling per bag) containing 40 L of locally sourced sandy loam Menangle soil (see Drake *et al.*, 2015 for details on soil characteristics) with a field capacity of approximately 17 % volumetric water content (VWC). The soil was placed on a 5 L base layer of 20 mm blue metal (quarried crushed, aggregate rock) to promote drainage.

The experiment had a factorial design with two populations and two water treatments (BOO-W, BOO-D, HRI-W and HRI-D). Thirty-two saplings from each population were allocated to each water treatment. The two water treatments were well-watered (W), representing 100 % VWC of field capacity, and chronic water deficit (D), representing 50 % VWC of field capacity (maintained within 43-57 % of saturated VWC field capacity).

An automated soil moisture sensor-triggered drip irrigation system was used to maintain soil moisture at the treatment set conditions. The irrigation system contained 49 two-wire TDT soil moisture sensors (Acclima Inc, ID, US) wired to two CS3500 2-wire controllers (Acclima Inc, ID, US). TDT sensors were inserted into target plant bags with up to three additional saplings from the same population and water treatment plumbed to these target saplings, thereby forming an irrigation zone of four saplings. The three saplings in the irrigation zone that did not contain sensors were assumed to have a VWC level similar to the sapling with the sensor. All saplings in an irrigation zone were irrigated according to soil moisture levels detected by the TDT moisture sensor in the target plant bag based on hourly readings.

Saplings were grown under well-watered conditions in the *establishment phase* from 30 August 2017-17 November 2017 (80 days). One cup of 1:300 diluted (as directed) liquid fertiliser was added monthly during this phase (N: P: K ratio of 24:3.5:16 with trace elements, all-purpose water-soluble fertiliser, Debco, VIC, Australia).

### Treatment phase

Water treatments were commenced on 18 November 2017. After 4 months of growth under water treatments (122 days), a subset of five replicate saplings per population × treatment combination, were used to estimate specific leaf area (SLA), minimum stomatal conductance (g_min_), branch capacitance (*C*) and pressure-volume curves (PV curves; see below for measurement details). An additional 11-18 replicate saplings per population × treatment combination were watered to field capacity prior to measurement of percent loss of conductivity (PLC) curves (see below). Slow release fertiliser (15 g Scotts Osmocote^®^ Plus Trace Elements: Native Gardens) was added monthly during this phase.

### Drought hardening phase

The remaining eight replicate saplings from each population × treatment combination containing TDT soil moisture sensors were allocated to measurements of time to hydraulic failure (THF) after almost 6 months under treatments (174 days). All THF saplings were drought-hardened by withholding water until individual saplings reached TLP (determined from PV curves). Pre-dawn leaf water potentials (Ψ_pd_) were measured using a Scholander-type pressure chamber (PMS Instruments, Corvallis, OR, USA) to confirm that every sapling achieved TLP. Saplings were subsequently watered to field capacity and allowed to recover for three days with Ψ_pd_ measured in a subset of saplings from each population × treatment combination to confirm rehydration.

### Drought to hydraulic failure phase

Water was then withheld from all THF saplings starting June 4 2018. Leaf Ψ was measured regularly during the final dry-down phase to determine when saplings reached the Ψ associated with 90 % stomatal closure (*P*_*gs90*_) and *P*_88_. For smaller saplings, leaves for Ψ measurements were alternately taken from similar sized saplings from the same population × treatment to minimise impacts of leaf removal on desiccation time.

Once individual saplings attained or exceeded *P*_88_ the whole sapling was cut at the soil surface, and leaves and stems were separated and weighed immediately using a balance. Leaves and stems were dried for at least 48 hours at 70 °C prior to dry mass determination.

Air temperature, relative humidity (Rotronic HygroClip2, HC2-S3, Rotronic Intruments Corp, NY, USA) and photosynthetically active radiation (Apogee SQ-420 PPFD sensor, Apogee Instruments Inc., UT, USA) were monitored during the experiment at 30 minute intervals at two locations within the poly-tunnel using a Campbell logger (Fig. S1; CR1000, Campbell Scientific, QLD, Australia). Conditions inside the poly-tunnel were monitored throughout the experiment, and side curtains were opened or closed to modify air circulation.

Air temperature and VPD inside the poly-tunnel varied substantially across seasons during the 11-month duration of the experiment (Fig. S1). Average temperature and VPD were 21°C and 1.16 kPa, respectively. During the *drought to hydraulic failure phase*, mean air temperature and VPD were 12.2 °C and 0.6 kPa, respectively. During the treatment phase, W saplings had an average of 16.9 % VWC, while D saplings had 7.3 % VWC (Fig. 1).

**Figure 1.**
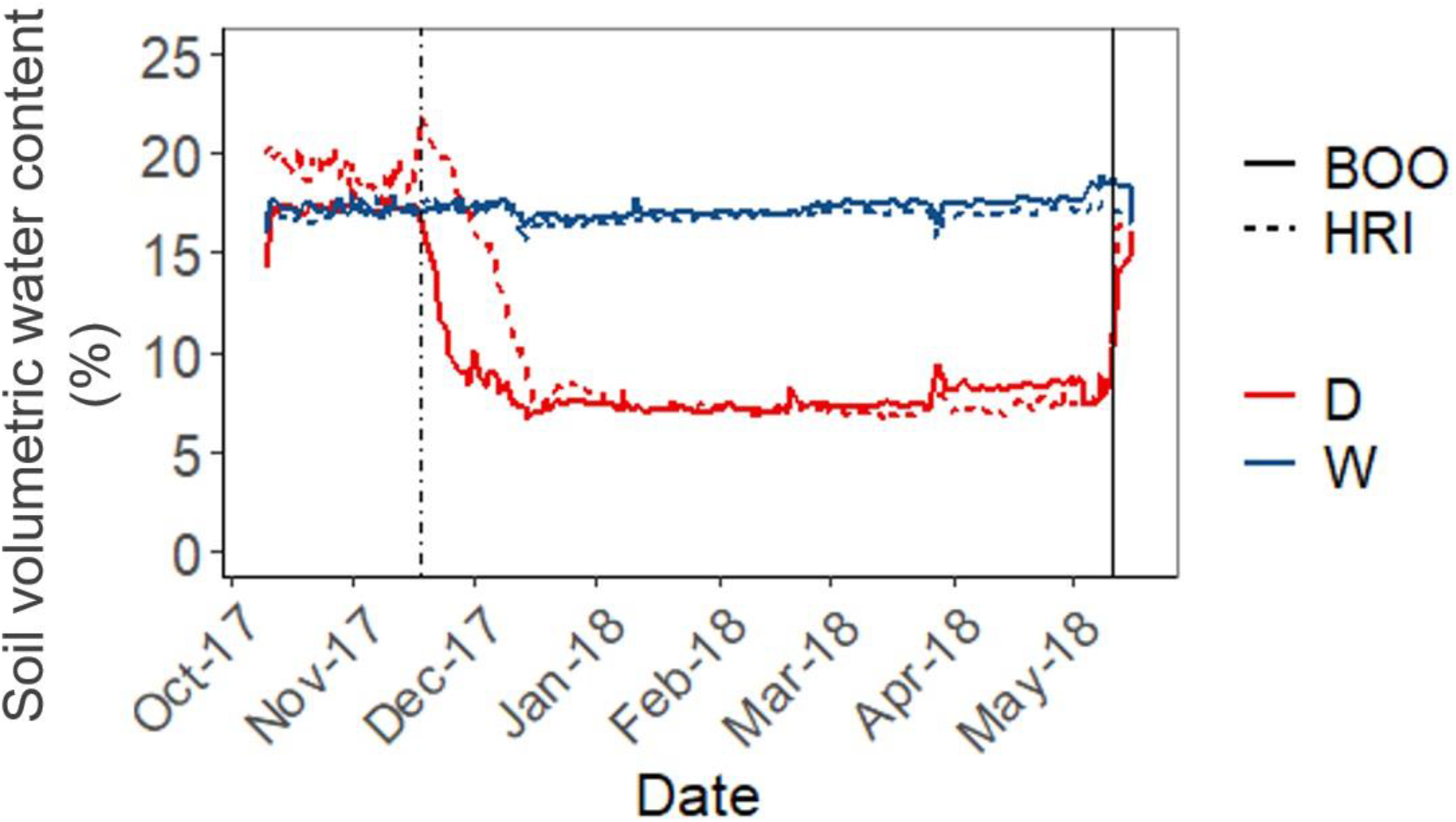
Mean daily soil volumetric water content (%) available to saplings in cool-wet BOO (solid line) and warm-dry HRI (broken line) populations under well-watered (W, blue) and water deficit (D, red) treatments. The broken vertical line indicates the date for commencement of the water treatments. The solid vertical line indicates rewatering of all trees and the commencement of the drought to hydraulic failure.

### Growth, leaf area and total water storage

Growth was measured in all saplings monthly from 7 September 2017 until 12 March 2018. Stem basal area was calculated from two perpendicular diameter measurements using digital callipers (mm) 2 cm above the soil surface. Sapling height (h, cm) was measured and stem volume calculated as:

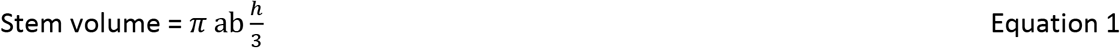

where ‘a’ and ‘b’ are perpendicular radii of the stem.

Leaf area was estimated (referred to as ‘estimated leaf area’) for THF saplings monthly from 13 November 2017 until 12 March 2018. Five leaves per sapling were selected and leaf area was measured independently for each using a 1 cm^2^ grid. All leaves on the sapling were counted and total sapling leaf area was estimated from the leaf number multiplied by the average leaf area for the five representative leaves.

Total projected leaf area (*A*_L_) was calculated in THF saplings when saplings were harvested. Values of *A*_L_ were calculated for individuals by multiplying the mean population × treatment specific leaf area (SLA, see supplementary materials) by total leaf dry mass.

The maximum amount of water in the aboveground biomass (*V*_w_ (g)) was calculated for each individual THF sapling at the end of the *time to hydraulic failure phase* by multiplying aboveground (leaves and stems) dry mass by the mean saturated water content of shoots calculated for each population × treatment.

### Pressure-volume analysis

A few days prior to the end of the *treatment phase*, one recently-matured leaf per sapling was sampled at pre-dawn from five replicates per population × treatment. PV curves were conducted according to Maréchaux et al. (2015; see supplementary materials for measurement details). PV traits were estimated according to Lenz, Wright, and Westoby (2006). Relative leaf water content and 1/Ψ_leaf_ were plotted and TLP was determined as the point where the line became non-linear.

### Stomatal closure

The leaf Ψ associated with stomatal closure (*P*_*gs90*_) was determined as 90 % loss of stomatal conductance (*g*_*s*_) in response to decreasing soil moisture based on stomatal closure curves measured for each population × treatment combination during the *treatment phase*. These measurements were conducted on a separate subset of 6-9 replicate saplings. Saplings were fully hydrated and leaf Ψ_pd_ measurements were taken on a recently mature leaf using a pressure chamber. Maximum *g*_*s*_ was measured in the same individuals on an adjacent leaf between 8:30 AM and 12:30 PM using a Licor-6400XT (Licor Inc., Lincoln, NE, USA) at saturating light (1500 μmolm^−2^s^−1^), ambient CO_2_ (400 μl l^−1^), relative humidity 50– 80%, and block temperature set to the maximum temperature forecasted on the measurement day. Water was then withheld from these saplings and periodically paired measurements of leaf Ψ_pd_ and *g*_*s*_ were measured until they reached *P*_*gs90*_. When plants had reached *P*_*gs90*_, *g*_*s*_ and Ψ_pd_ were measured on two consecutive days, to confirm that stomata had closed in response to soil water deficit, saplings were subsequently rewatered to ensure full recovery.

### Shoot hydraulic capacitance

Five replicates from each population × treatment were allocated to aboveground shoot (leaves and stems) capacitance (*C*) measurements. From 19 March 2018, following 4 months of growth under water treatments, shoots were cut at the stem base under water at pre-dawn. Shoots were transported to the laboratory and allowed to desiccate, during which time paired measurements of total shoot mass and Ψ_leaf_ were taken (see supporting materials for more details).

Relative water content (RWC) of the shoot was plotted against Ψ_stem_ for each population × treatment and an exponential curve was fitted to the portion of the curve following *P*_*gs90*_ (see Fig. S3). RWC at *P*_*gs90*_ (θ_0_) and *P*_88_ were calculated from the equation of these curves for each population × treatment by substituting *P*_*gs*90_ and *P*_88_ as *x* into the equation to get the RWC at *P*_*gs*90_ and *P*_88_ (*y*). The difference in RWC content between these two points (here after referred to as “Δ RWC”) was calculated as:

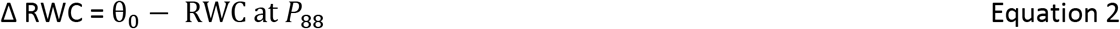

Shoot *C* was calculated from the linear portion of the post-turgor loss relationship between RWC and Ψ (g g^−1^ MPa^−1^).

### Stem hydraulic vulnerability curves

Percent loss of conductivity curves (PLC) were estimated from 11-18 replicate saplings per population × treatment combination. The number of saplings required to create a PLC curve was dependent on the size of the saplings. Sampling for PLC measurements was conducted over March-July 2018 by slowly desiccating subsets of saplings *in situ* in their pots (Tyree *et al.*, 1992). Saplings were dried to a target Ψ_stem_ to populate the PLC curves and then either a branch or the whole sapling (maximum vessel length varied from 14 – 74.5 cm) was excised to measure stem PLC.

Percent loss of conductivity was measured using a flow meter (Liqui-Flow L10, Bronkhorst High-Tech BV, Ruurlo, Gelderland, The Netherlands) and analysed using the FlowDDE and FlowPlot software (Version 4.76 and 3.34, respectively, Bronkhorst, FlowWare). Initial flow rates (*K*_*init*_) were measured, stem segments were then flushed with 2 mmol KCl solution for 30 minutes (1 bar pressure) and then maximum flow rate (*K*_*max*_) was measured. PLC was calculated as:

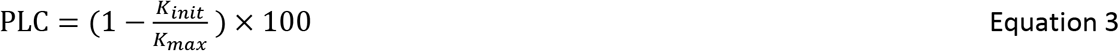

See supporting materials for further details of methods

### Minimum leaf conductance

Minimum leaf conductance (g_min_) was measured according to the mass loss of water from detached leaves (Duursma *et al.*, 2019). Two recent fully expanded leaves were sampled at pre-dawn from five replicate saplings per population × treatment. Leaves were scanned for leaf area and weighed immediately using a balance (0.0001 g). Samples were slowly desiccated in a growth chamber and paired measurements of leaf mass and measurement time were recorded (see supplementary materials for further method details).

Changes in leaf mass with time had an initial exponential decay relationship with high water loss prior to *P*_*gs90*_. Minimum leaf conductance (g_min_) was calculated from the slope of the latter linear region of the relationship between decreasing leaf mass (g) and increasing time (mins). This was converted from g^−1^ min^−1^ to mmol m^−2^ s^−1^ by dividing by projected leaf area (m^−2^) and the mean chamber VPD, (approx. 0.5 kPa), and converting the mass loss from g to mmol H_2_O.

### Time to hydraulic failure

Time to hydraulic failure (THF, kPa hr) was calculated as the sum of hourly VPD between the hour corresponding to leaf Ψ_pd_ on the day of *P*_*gs90*_ and the hour corresponding to *P*_88_. THF was standardised based on the relative cumulative VPD hours (mean daily VPD × 24 hours + previous day’s VPD hours with time zero at *P*_*gs90*_) across all saplings in each population × treatment over the course of the final dry-down phase.

### Statistical analysis

PLC and *P*_*gs90*_ curves were analysed by fitting sigmoidal or Weibull curves for each population × treatment group using the *fitplc* function in the fitplc R package (Duursma & Choat 2017). *P*_88_ and *P*_50_ were calculated as the Ψ resulting in 88 % and 50 % loss of conductivity from the PLC curves, respectively. *P*_*gs90*_ was calculated as the Ψ resulting in 90 % loss of *g*_*s*_ from the maximum values. *P*_*gs90*_ did not differ significantly between treatments within populations, therefore treatments were pooled within populations and curves. *P*_88_, *P*_50_ and *P*_*gs90*_ were obtained from the curves using the *getPx* function. Significant differences for *P*_88_ between population × treatment combinations and populations for *P*_*gs90*_ were determined from non-overlapping 95 % confidence intervals.

Significant differences amongst G×E interactions were tested using a generalised linear model using the *glm* function with a Gaussian family with treatment and genotypes as interacting variables. *Post-hoc* Tukey tests (*glht* function) were performed to identify differences among treatment × population combinations at an α level of 0.05. To test for G×E interactions for RWC traits and THF, linear models were used followed by the *post-hoc* Tukey test (*glht* function). For the THF linear model, cumulative VPD and associated Ψ_stem_ during the sapling final dry down phase from *P*_*gs*90_ to *P*_88_ for all saplings was used with cumulative VPD as the response variable, Ψ_stem_ as the explanatory variable and treatment × population as the covariate. For RWC traits (θ_0_ and RWC *P*_88_), whole sapling RWC and associated Ψ_stem_ from *C* curves were used with whole sapling RWC as the response variable, Ψ_stem_ as the explanatory variable and treatment × population as the covariate. Normality and homogeneity of variance were tested and data were log transformed, when necessary.

Mean annual temperature (MAT) and precipitation (MAP), mean maximum temperature of the warmest month (MaxT) and precipitation of the driest month (PreDM) were extrapolated from WorldClim data sets (www.worldclim.org) and selected to characterise population climate-origin. Aridity index (AI; mean annual precipitation/mean annual evapotranspiration) was obtained from the raster downloaded from CGIAR-CSI (http://www.cgiar-csi.org/) and 1/AI was used.

All analyses were conducted in R v1.2.1335 (R Development Core Team, 2018).

## Results

### Plant growth under the watering treatments

Prior to the commencement of the water treatments, populations did not differ significantly in stem volume or estimated leaf area (estimated leaf area *P* = 0.487, stem volume *P* = 0.705). Stem volume differed significantly among treatments after 23 days of treatment (*P* < 0.0001), and estimated leaf area showed significant differences after 59 days (*P* < 0.0001). At the end of the treatment phase (174 days of treatment), plant size was significantly different among treatments (estimated leaf area *P* < 0.0001, stem volume *P* < 0.0001). Stem volume of W saplings was 2.6 times larger than D saplings, and estimated leaf area 2.2 times greater in W saplings than D saplings when populations were pooled (Fig. 2). Mean stem volume and estimated leaf area did not differ between populations grown under the W treatment, but significant differences in stem volume (*P* = 0.001) and estimated leaf area (*P* = 0.002) were observed between populations grown under the D treatment, with the smallest leaf area and lowest stem volume in the HRI population. At the end of the treatment phase, estimated leaf area and stem volume had a significant G×E interaction between treatment and population (estimated leaf area G×E *P* = 0.013, stem volume G×E *P* = 0.01).

**Figure 2.**
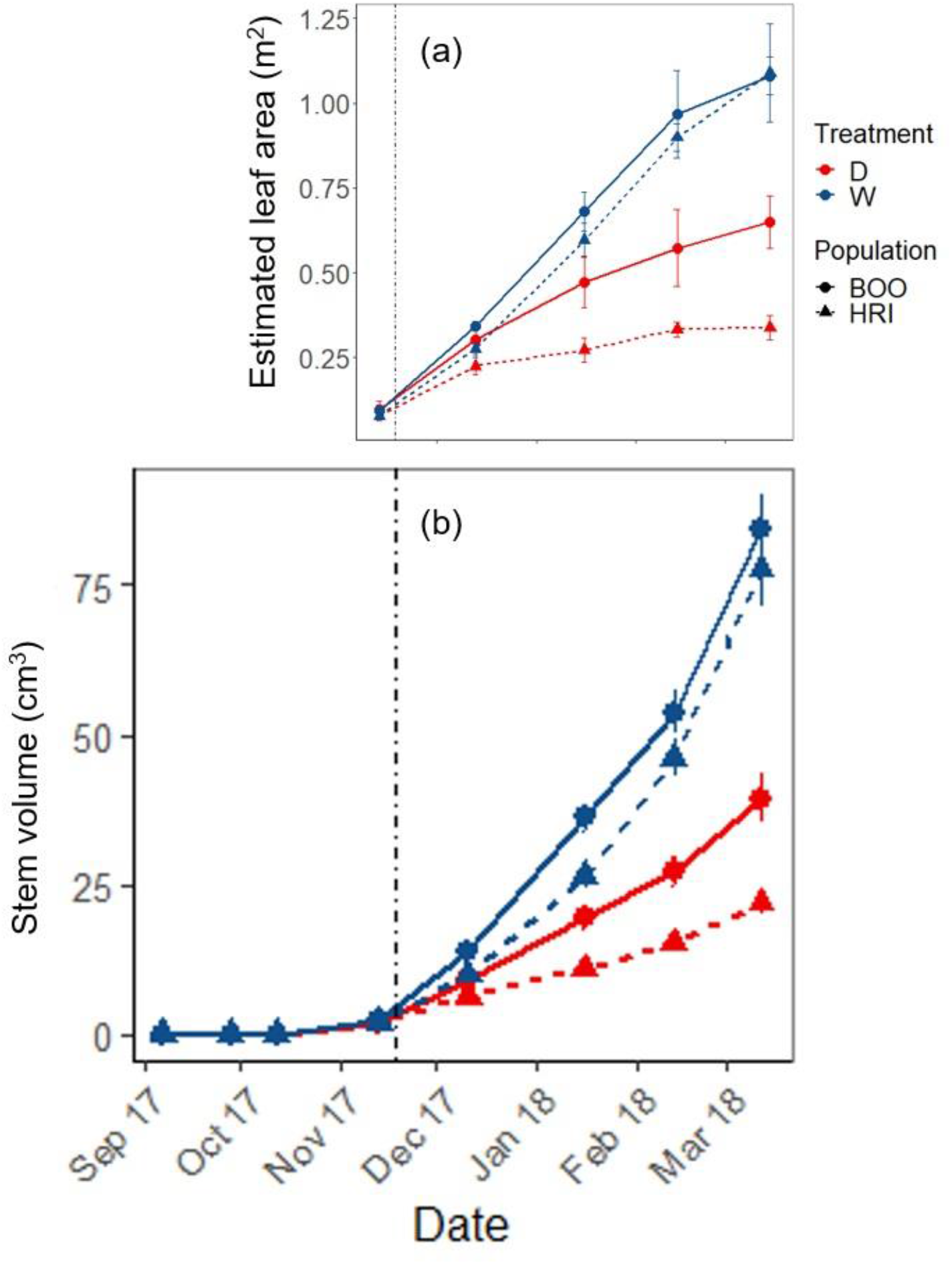
Estimated whole plant leaf area (m^2^, a) and stem volume (cm^3^, b) over time in *C. calophylla* saplings from the cool-wet population BOO (circles, solid lines) and warm-dry population HRI (triangles, dashed lines) saplings grown under soil water deficit (red, D) and well watered (blue, W) treatments. Vertical broken line indicates commencement of water treatments.

### Hydraulic and allometric traits

The D treatment had a significant effect on both pre-dawn (*P* < 0.001) and midday (*P* = 0.003) leaf water potentials relative to the W treatment, but there was no genotypic variation detected (pre-dawn and midday *P* > 0.05) in saplings measured after 122 days under water treatments in the *treatment phase*.

Traits showing a G×E interaction included *P*_88_, Δ RWC and the ratio of total stored shoot water and leaf area (*V*_w_/*A*_L_). The effect of the D treatment on *P*_88_ was significantly stronger in the warm-dry population HRI than in the cool-wet population BOO (Table 2; Fig. **3c**). *P*_88_ was significantly greater (more negative) for HRI-D saplings (8.35 -MPa) than for other treatment × populations, which were less negative and therefore less drought tolerant (ranging from 6.24 to 6.46 -MPa; Table 2; Fig. 3; Fig. S5). *P*_50_ was highest in HRI-D saplings, and although it was not significantly different to the HRI-W saplings, it was significantly greater compared to the BOO-D and BOO-W saplings (Table 2).

**Table 2.**
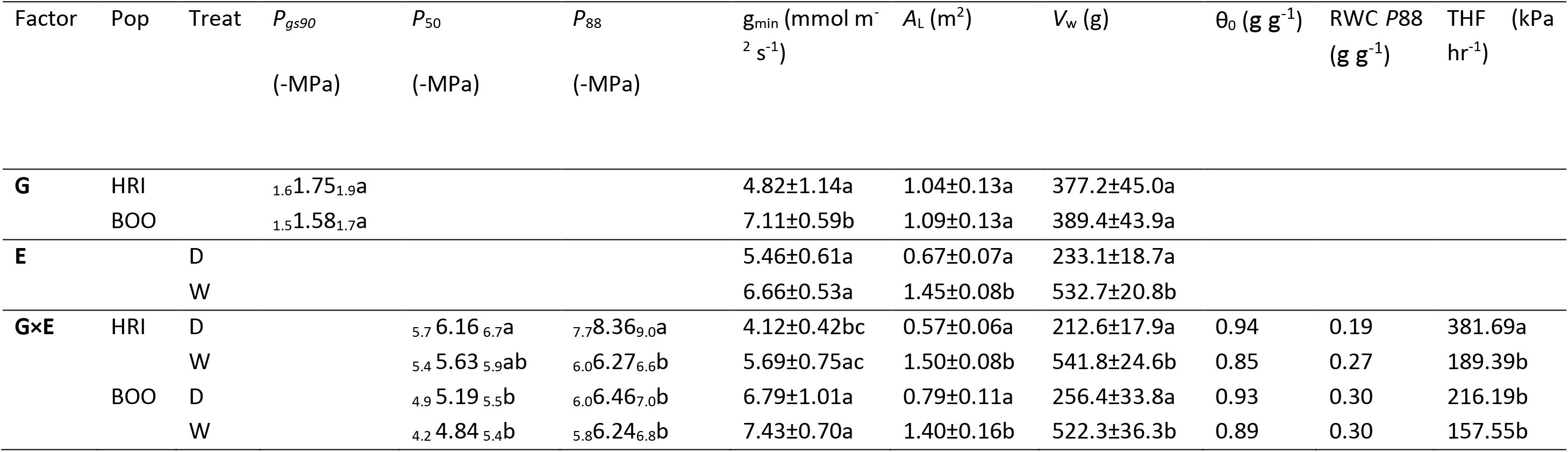
Mean in hydraulic and allocation traits and time to critical failure (THF, VPD (kPa) hours). Values are first averaged for HRI and BOO populations (Pop, G), then for D and W treatments (Treat, E) and then given for each genotype × environment interaction (G×E). Plant traits are *P*_*gs90*_ (leaf water potential at stomatal closure, -MPa), *P*_50_ (water potential at 50 % loss of conductivity, -MPa), *P*_88_ (water potential at 88 % loss of conductivity, -MPa), g_min_ (minimum leaf conductance, mmol m^−2^s^−1^), *A*_L_ (projected total leaf area, m^2^), θ_0_ (relative water content at stomatal closure, g g^−1^), RWC *P*_88_ (relative water content at *P*_88_, g g^−1^), *V*_w_ (total amount of stored water, g). Significant differences for *P*_*gs90*_, *P*_50_ and *P*_88_ are determined from non-overlapping 95% confidence intervals which are presented in subscript. For other parameters, ±1SE are shown. Significant differences are indicated by letters.

**Figure 3.**
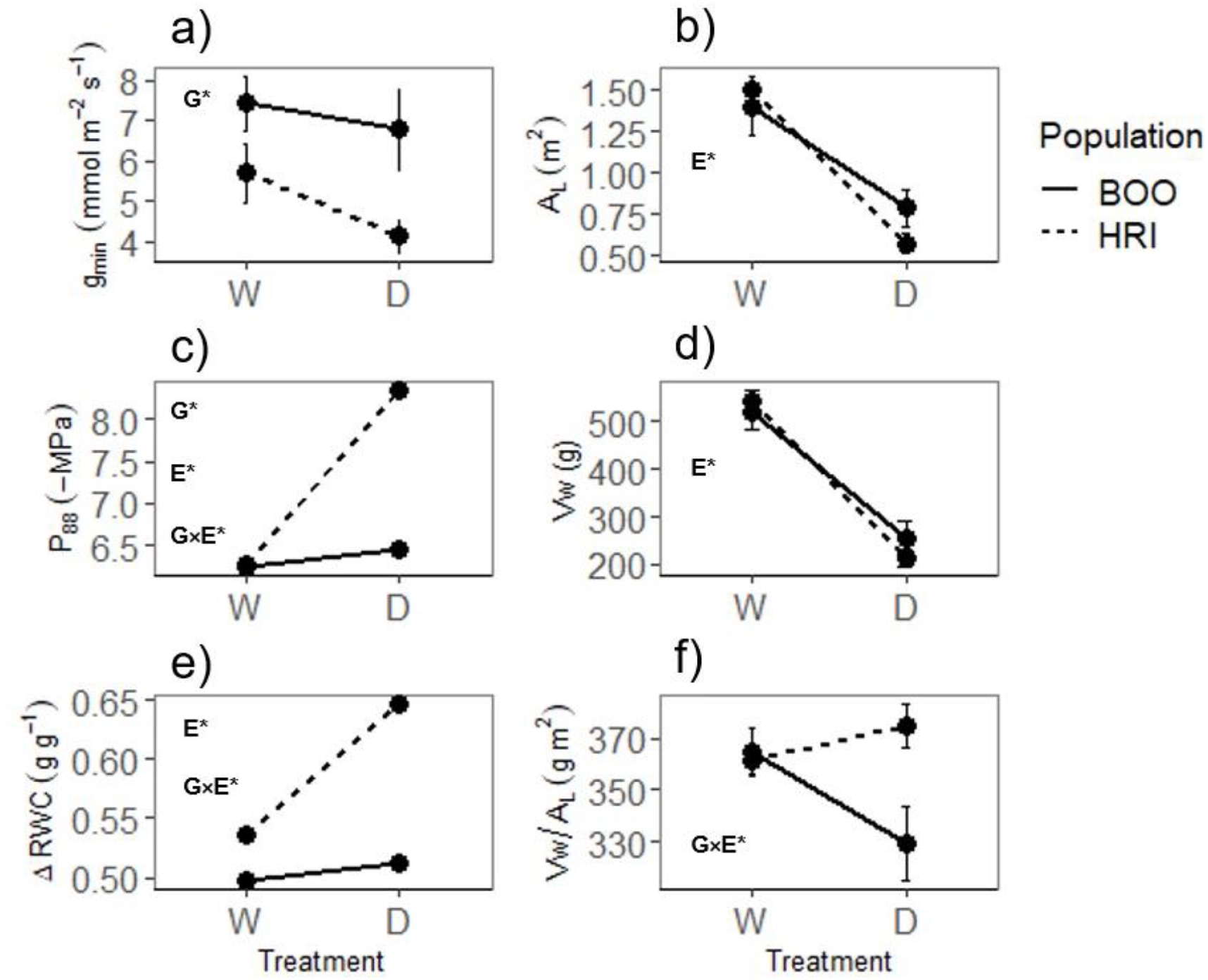
Reaction norm plots indicating degree of plasticity in traits across well-watered (W) and 50 % of field capacity (D) treatments for HRI (dashed lines) and BOO (solid lines) populations. Traits presented: (a) g_min_ (minimum leaf conductance, mmol m^−2^ s^−1^), (b) *A*_L_ (whole plant projected leaf area, m^2^), (c) *P*_88_ (water potential at 88 % loss of conductivity, -MPa), (d) *V*_w_ (total plant stored water, g), (e) ΔRWC (change is relative water content between stomatal closure and *P*_88_), and f) *V*_w_/A_L_ (ratio of total stored water and total plant leaf area, g m^2^). Significant genotype (G), environment (E) and G×E interactions are indicated by letters and asterisks. Error bars show ± 1 SE.

For Δ RWC, the D treatment had a greater impact on HRI saplings than BOO saplings and there was a significant G×E interaction (Fig. **3e**). For *V*_w_ and *A*_L_ traits there was no significant G×E interaction, but when these traits were combined as *V*_w_/*A*_L_ there was a significant G×E interaction with lower values recorded in HRI-D saplings (*P* = 0.023; slopes *P* = 0.018, intercepts *P* = 0.018; Fig. **3b, d**). *A*_L_ and *V*_w_ traits did not differ significantly between populations (*A*_L_ *P* = 0.775, *V*_w_ *P* = 0.848; Fig. 3). However, there was a significant E effect detected in these traits. Saplings grown under the D treatment had a significantly smaller *A*_L_ and lower *V*_w_ than W treated saplings (*A*_L_ *P* < 0.0001, *V*_w_ *P* < 0.0001; Fig. **3b, d**). These traits were significantly correlated (r^2^ values D saplings 0.94; W saplings 0.90; *P* < 0.0001; Fig. S4). For both *A*_L_ and *V*_w_, there were shared slopes and intercepts across treatments and populations.

One trait, g_min_, showed a significant G effect but no E effect. There was a trend of lower g_min_ in D treated saplings relative to W saplings, but this was marginally not significant (*P* = 0.056). HRI saplings exhibited significantly lower g_min_ than BOO saplings (*P* = 0.004; Fig. **3a**). g_min_ was lowest in HRI-D saplings and greatest in BOO-W saplings with no G×E interaction for this trait. *P*_*gs90*_ was higher in HRI saplings (1.75 -MPa) relative to BOO saplings (1.58 -MPa), but was not significant (treatments pooled). Similarly, the range of θ_0_ was small (0.85 – 0.94 g g^−1^). HRI-D saplings had the lowest RWC at *P*_88_ at 0.19 g g^−1^, while other treatment × populations had higher values ranging from 0.27 - 0.30 g g^−1^ (Table 2).

There were no significant differences detected for TLP or SLA (mean SLA = 6.3 m^2^ kg^−1^ and mean TLP −2.82 MPa; *P* > 0.05). Mean *C* was 0.237 g g^−1^ MPa^−1^ and no significant differences were found for this trait (*P* > 0.05; Fig. S3).

### Time to hydraulic failure

A G×E interaction was detected in THF saplings as HRI-D saplings had significantly longer dry-down times than other treatment × populations (*P* < 0.05). The D treatment substantially increased THF in HRI but not in BOO saplings, and HRI-D saplings had a substantially longer THF than all other saplings (381.69 VPD hrs; Table 2; Fig. 4). HRI-W, BOO-D and BOO-W saplings exhibited similar THF values (189.39, 216.19 and 157.55 VPD hrs, respectively). THF for HRI-D saplings was 202 % greater than HRI-W saplings, and BOO-D was 137 % higher than BOO-W.

**Figure 4.**
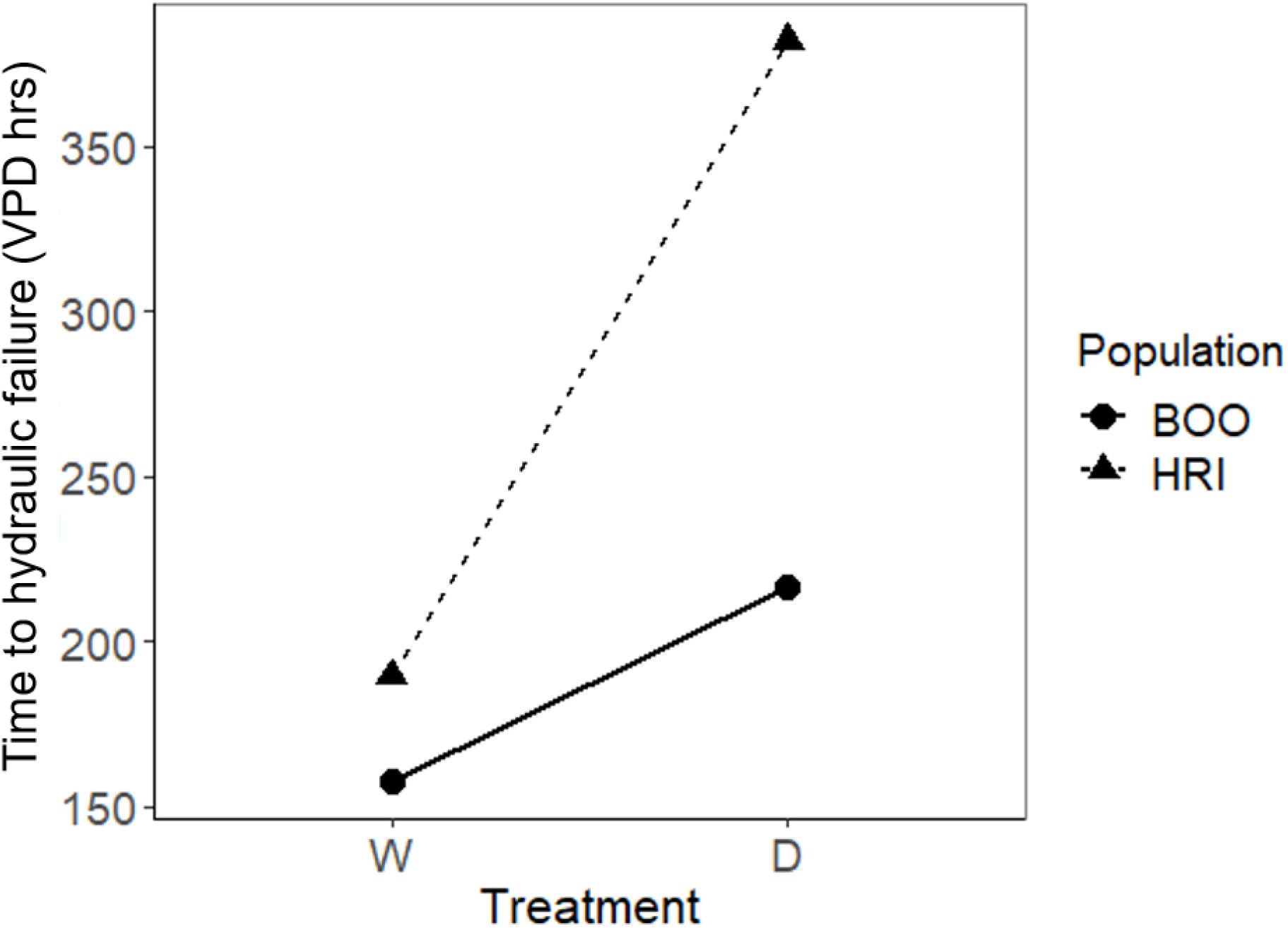
Time to hydraulic failure (VPD hours) in Boorara (BOO, circles, solid line) and Hill River (HRI, triangle, broken line) populations grown under well-watered (W) and water deficit (D) treatments.

## Discussion

Time to hydraulic failure (THF) in *C. calophylla* saplings from two differentiated populations grown under well-watered (W) and water deficit (D) treatments showed a significant G×E effect, indicating different capacities to respond to climate change-induced drought among populations. Under the D treatment, the hot-dry climate population (HRI) increased THF more than the cool-wet climate population (BOO), likely because of the capacity of the HRI population to increase *P*_88_, while decreasing the rate of residual water loss (g_min_). A significant G×E interaction was observed in THF in that HRI-D saplings had higher values of THF relative to other population × treatment combinations, which supported hypothesis 1. Observation of genotypic and environmental variation occurred in THF, which supported hypotheses 2 and 3.

### Hydraulic and allocation traits influencing time to hydraulic failure

Numerous hydraulic and allometric traits likely influenced the THF. In *C. calophylla* saplings, THF and *P*_88_ expressed similar G×E trends with the longest THF and highest *P*_88_ (greater drought tolerance) in HRI-D saplings and similarly low values for other population × treatment combinations. Thus, *P*_88_ was likely to have a large influence on THF in these saplings. Although there is still debate regarding the exact mechanisms of drought-induced mortality (Hammond *et al.*, 2019), the water potential at stem *P*_88_ provides a useful measure of critical (but not necessarily lethal) hydraulic dysfunction, with recovery typically very slow (Kursar *et al.*, 2009; Urli *et al.*, 2013). The significantly greater *P*_88_ in HRI-D saplings indicates a capacity to tolerate a greater level of water stress during drought in comparison with saplings from other population × treatment combinations. In support of these findings, Blackman et al. (2019b) found a strong relationship between *P*_50_ and the time for saplings to desiccate from stomatal closure (*P*_*gs90*_) to *P*_88_ (*t*_*crit*_) in eight *Eucalyptus* species, with those exhibiting greater stem *P*_50_ expressing a longer *t*_*crit*_. Additionally, Blackman et al. (2019a) found the stomatal-hydraulic safety margin (*P*_*gs90*_ - *P*_50_) was more influential in determining desiccation time in three angiosperm and one gymnosperm tree species than *P*_50_ and *P*_88_ traits. *P*_50_ and *P*_88_ were significant predictors of tree mortality anomalies across species in a global meta-analysis (Anderegg *et al.*, 2016). These cavitation resistance traits are therefore highly informative for understanding drought resistance in plants, mortality thresholds and desiccation time.

g_min_ was lowest in HRI-D saplings, although only significantly lower than BOO-D and BOO-W saplings. Low g_min_ contributes to slow plant water loss after *P*_*gs90*_, thereby influencing plant desiccation time (Burghardt & Riederer 2003; Brodribb *et al.*, 2014) and would have played a role in increasing THF in HRI-D saplings through a reduced rate of water loss. g_min_ was an important trait differentiating between desiccation time to critical levels in gymnosperms and angiosperms in the rain-out shelter experiment by Blackman et al. (2019a). When g_min_ is expressed on a whole plant leaf area basis, it provides a measure of the whole plant residual water loss rate. In the present study, *C. calophylla* saplings grown under the D treatment exhibited significantly lower *A*_L_ compared to W saplings. HRI-D saplings had a small *A*_L_ and low g_min_, thereby reducing the rate of plant residual water loss and likely increasing THF in these saplings. Total leaf area is an important plant trait strongly influencing plant water loss under drought conditions (Choat *et al.*, 2018) and large trees with a large leaf area are likely to die sooner than those with a smaller leaf area (McDowell & Allen 2015). Plants with the capacity to reduce *A*_L_ under drought conditions, as observed here, confers an advantage through more conservative water use strategy.

The ratio of total water content and canopy leaf area (*V*_w_/*A*_L_) did not show a trend suggestive of contributing to longer THF in HRI-D saplings relative to other population × treatment combinations. *V*_*w*_ provides a reserve for prolonging plant survival following *P*_*gs90*_ when plants are no longer extracting soil water. D treated saplings had significantly lower *V*_w_ compared to W saplings. However, the THF in HRI-D saplings was longer than HRI-W saplings, indicating that *A*_L_ is counteracting *V*_w_ in these saplings and *P*_88_ and g_min_ traits are likely to be more influential on THF. In contrast with these findings, Blackman et al. (2019b) found *V*_w_/*A*_L_ was highly significant in influencing *t*_*crit*_ in *Eucalyptus* species, suggesting that *V*_w_/*A*_L_ may be important among species, but less so within species.

### Genetic adaptation and Phenotypic plasticity

The hydraulic failure trait, *P*_88_ likely had the greatest influence on THF across treatments and populations. A significant G×E interaction was found for *P*_88_ with significant plasticity observed in the HRI population with greater drought tolerance in HRI-D saplings, but limited plasticity in the BOO population. Multiple interspecific comparison studies have revealed a strong correlation between *P*_50_ and precipitation at the climate-of-origin (Choat *et al.*, 2012; Dória *et al.*, 2018; Li *et al.*, 2018), suggesting an influence of adaptation on drought tolerance at the species level. However, studies relating intraspecific variation in cavitation traits with climate-of-origin are less common. In a previous *C. calophylla* population glasshouse study, *P*_50leaf_ was adaptive and related to temperature at the site-of-origin but not rainfall, and warm origin populations, including the HRI population, were more drought tolerant (more negative *P*_50leaf_) than cool origin populations (Blackman *et al.*, 2017). Across 10 provenances of European beech trees grown in a single field common garden site, *P*_88_ varied significantly across provenances but was not associated with aridity at site-of-origin (Hajek et al. 2016). Lamy et al. (2014) observed minimal plasticity and adaptive variation in *P*_50_ across six *Pinus pinaster* populations grown in a dry and wet common garden with rainfall reflecting mean annual precipitation experienced by the driest population and moderately wet populations. The common gardens experienced similar temperatures (Lamy *et al.*, 2014). Similar to the present study, Lopez et al. (2013) found a significant G×E interaction in *P*_88_ in *Pinus canariensis* with greater drought tolerance in hot, dry climate populations than mesic populations in a dry common garden, but not in a mesic common garden site. Combined, these findings suggest that variability in cavitation resistance may only be observed in populations at the species climate margins grown under water limited environments. Cavitation resistance traits are important for understanding tree mortality risk under climate change and additional studies that investigate genotype-by-environment interactions in these traits are necessary.

HRI-D saplings exhibited low g_min_, low RWC at *P*_88_, small *A*_L_ and a large Δ RWC contributing to greater drought tolerance that likely increased THF relative to other population × treatment saplings. A recent review of g_min_ (Duursma *et al.*, 2019) suggests that growth conditions influences g_min_ more than climate-of-origin; yet our study found genotypic variation in g_min_. For trees from dry environments, low g_min_ will provide an advantage under water deficit as the rate of plant water loss is reduced. Blackman et al. (2019b) did not observe systematic variation in g_min_ across *Eucalyptus* species from contrasting climates grown under common conditions. Few studies have investigated g_min_ variability across populations and water treatments, and our findings suggest further investigation is needed to improve our understanding of the contribution of this trait to plant desiccation time (Martin-StPaul *et al.*, 2017). While *A*_L_ measured at the end of the *drought to hydraulic failure phase* exhibited a treatment but not population effect, estimated leaf area measured over time during the *treatment phase* exhibited a significant G×E effect with significantly smaller estimated leaf area in HRI-D saplings relative to BOO-D saplings later in the *treatment phase*. Hajek et al. (2016) found significant genotypic variation in mean leaf size across 10 provenances of European beech and Pita et al. (2003) observed a G and E effect on leaf area across clones of *Eucalyptus globulus* grown under drought and well-watered conditions with a smaller leaf area recorded under the drought treatment. Leaf area is an important trait influencing tree desiccation time and we recommend its inclusion in future studies on time to critical failure.

The HRI population is situated at the warm and dry-end limits of the *C. calophylla* distribution and under climate change will likely experience higher temperatures and reduced rainfall in the future compared to its contemporary climate. A recent meta-analysis comparing tree mortality occurrences between dry-edge and range core populations indicates mortality was more prevalent at the dry-edge margins relative to the range core and that drought tolerance mechanisms did not effectively buffer against drought-induced mortality (Anderegg *et al.*, 2019). Therefore, the significant capacity for the HRI *C. calophylla* population to adjust cavitation traits under contrasting water availability leading to increased drought tolerance may not sufficiently buffer against global change-type droughts. Despite the lower capacity for the cool, wet BOO population to adjust cavitation traits through phenotypic plasticity, it may experience lower mortality rates than the HRI population because drought events in the region are milder.

### Limitations and field growing trees

This study provides insight into intraspecific and plastic variation in THF in saplings, building on recent studies that examine THF across species (Blackman *et al.*, 2019a, b). The sample of intraspecific variation was limited to two populations in the present study; however, these populations were selected because they exhibited genetic differentiation (Ahrens *et al.*, 2019b) and were situated at the geographical extremes of *C. calophylla’s* distribution, thereby experiencing contrasting temperature and rainfall regimes.

We acknowledge that this study has limitations with extrapolation to trees growing in the field. The large pots enabled accurate control of soil water availability, but maintenance of drought treatments in the field is challenging. Like many Mediterranean-type ecosystem woody species, *C. calophylla* has a deep rooting system (Nardini *et al.*, 2014) and this may contribute to total tree water storage and capacitance (*C*) and hence, may prevent trees from experiencing significant water stress close to hydraulic failure due to access to deep sources of water. Blackman et al. (2019b) found an improvement to the *t*_crit_ model fit when root water storage was added to *V*_w_. Furthermore, as trees mature the expression of traits influencing THF may vary as allometric relationships shift (Hartmann *et al.*, 2018). Large trees store a greater volume of water, have a larger projected leaf area and higher *C* than smaller trees (Scholz *et al.*, 2011). *C* may play a greater role in reducing cavitation through its buffering effect differentially influencing THF in larger trees relative to smaller trees (Scholz *et al.*, 2011). The effect of higher water loss as a result of a large total leaf area in large, mature trees may be counteracted by the corresponding large *V*_w_ and higher *C*. Cavitation resistance traits may remain relatively consistent across life stages; however, tall trees are more vulnerable to cavitation than shorter trees (Koch *et al.*, 2004).

*Corymbia calophylla* trees have the capacity to resprout after droughts and fires (Matusick *et al.*, 2016). Resprouting will have implications on tree mortality in the field in response to drought events as trees may experience dieback during extreme droughts and appear to be dead, but recover through resprouting. Resprouting tree species should be monitored after drought events occur to confirm mortality. The long-term implications of resprouting and dieback cycles on forest structure are unknown (Walden *et al.*, 2019).

## Conclusion

Tree mortality events in response to extreme weather are likely to become more common under predicted climate projections in south Western Australia and other Mediterranean-type ecosystems. In extreme drought years in this region, soil water reserves relied on by deep-rooted Mediterranean trees during hot summer periods will diminish and may result in tree mortality (Canadell *et al.*, 1996; Evans *et al.*, 2013; Matusick *et al.*, 2013). In the current study, we found significant intraspecific differences in the time for plants to desiccate from *P*_*gs90*_ to *P*_88_. These differences in THF were driven by adaptive plasticity in multiple drought response traits. There were several plant traits influencing drought-associated critical failure that are not often typically measured in tree drought studies (g_min_ and *A*_L_), while *P*_88_ was confirmed as a highly informative drought trait. Drought-associated hydraulic failure is complex, and our study provides evidence for significant G, E and G×E interactions driving trait differences related to drought tolerance and plant desiccation time in an ecologically important tree. Assisted gene migration may be a viable management option for the species, involving the introduction of genetic material from warm-dry populations into the cool-wet regions, thereby increasing tolerance in more vulnerable southern populations to the increasingly drying climate. Without management intervention, *C. calophylla* cool-wet climate populations with limited plasticity in key drought tolerance traits may be at risk of drought-induced hydraulic failure. These results illustrate that the adaptive capacity to cope with novel climate conditions may be naturally occurring within this tree species.

## Supporting information

see supplementary materials

## Acknowledgements

The authors would like to thank Katie Rolls, Aurelien Estarague, Renee Smith and Elisa Stefaniak for assistance with conducting measurements and Craig Barton and Burhan Amiji for poly-tunnel facility maintenance and set up. We would also like to thank Margaret Byrne, Richard Mazennec, Katinka Ruthrof and Giles Hardy. This research was funded by the ARC Linkage Project (LP150100936).

