## Supplementary material for "Adaptive plasticity in plant traits increases time to hydraulic failure under drought in a foundation tree": see supplementary materials

**Supplementary Information**

**Methods**

***Pressure volume curves***

A few days prior to the end of the *treatment phase*, one recently-matured leaf per sapling was sampled at pre-dawn from five replicates per population × treatment. Leaves were transported to the laboratory, in a sealed plastic bag inside an insulated box, where they were then transferred to a beaker containing water to immerse the petioles and rehydrated in a dark refrigerator for approximately 2 hours. Leaf Ψ and leaf fresh mass were immediately measured using a pressure chamber and a balance (0.001 g), respectively, and were then measured periodically as leaves desiccated slowly in the laboratory. Leaves were dried in an oven at 70 °C for at least 48 hours and weighed to obtain dry mass.

***Stem hydraulic capacitance***

Five replicates from each population × treatment were allocated to stem capacitance (*C*) measurements. From 19^th^ March 2018, following 4 months of growth under water treatments, whole sapling aboveground shoots (leaves and stems) were cut at the base of the stem under water at predawn. The cut surface was immediately sealed with parafilm and the shoots were covered with an opaque plastic bag and transported to a large refrigerator where the cut stems were immersed in water and shoots were rehydrated for two hours in the dark. The fully hydrated shoots were transported to a laboratory where whole shoot mass was measured immediately using a balance (0.01 g) and Ψ_stem_ was measured using a pressure chamber. Stem water potential was derived from covering the whole shoot with an opaque plastic bag 30 minutes prior to measuring Ψ_leaf_ thereby enabling leaf and stem Ψ to reach equilibrium. Shoots were allowed to slowly desiccate in the laboratory over the course of 3-4 days and shoot mass and Ψ_stem_ were measured periodically until the branch reached or exceeded *P*_88_. Leaves and stems were then separated and dried at 70 °C for at least 48 hours (in addition to Ψ leaves) and weighed to obtain dry mass.

***Stem vulnerability curves***

Eleven-eighteen replicate saplings per population × treatment were allocated to PLC curves. Sampling for PLC measurements was conducted over March-July 2018 by slowly desiccating subsets of saplings *in situ* in their pots (Tyree et al. 1992). Saplings were dried to a target Ψ_stem_ to populate the PLC curves and then either a branch or the whole sapling was excised to measure stem PLC; *Corymbia calophylla* tree xylem vessel length varies substantially from 14 – 74.5 cm.

When sampling was conducted during day light hours, Ψ_stem_ was obtained by covering targeted recently mature leaves with plastic wrap and then aluminium foil for 30 minutes prior to sampling of leaves; this was done to halt further water loss from leaves and enable Ψ_stem_ and Ψ_leaf_ to come into equilibrium. Leaf water potential was then measured using a pressure chamber.

Whole saplings were cut at the stem base under water. For branch excision, saplings were placed on their sides and the base of the branch was immersed in water and cut under water. The cut section was placed in water and then immediately covered in parafilm. Saplings were covered with an opaque bag and transported to the laboratory. A 6-10 cm stem segment was targeted for PLC measurements (method according to Sperry et al. 1988). The first cut was made upstream at least 30 cm from the target stem segment under water. Xylem tension was allowed to relax for at least 10 minutes prior to further cuts being made. Subsequent cuts were made slowly, alternating upstream and downstream from the target stem section over 30-40 minutes.

Percent loss of conductivity was measured using a flow meter (Liqui-Flow L10, Bronkhorst High-Tech BV, Ruurlo, Gelderland, The Netherlands) and analysed using the FlowDDE and FlowPlot software (Version 4.76 and 3.34, respectively, Bronkhorst, FlowWare). The stem segment was then flushed with 2 mmol KCl solution with a pressure of 1 bar for at least 30 minutes and then maximum flow rate (*K_max_*) was measured. PLC was calculated as:

$PLC=(1-\frac{Kinit}{Kmax} )\times100$	Equation 3

***Specific leaf area***

A single recently mature leaf was sampled from between 8-17 replicate saplings from each population × treatment for SLA. Leaves were placed in sealed plastic bags in an insulated box and transported to a laboratory. Leaves were immediately scanned for leaf area using a LI-3100C leaf area meter (Licor, Lincoln, NE, USA) and then dried in an oven for at least 48 hours at 70 °C and weighed for dry mass. Specific leaf area was calculated as the leaf area (m^2^) divided by leaf dry mass (kg).

***Minimum leaf conductance***

Minimum leaf conductance (g_min_) was measured according to the mass loss of water from detached leaves (Duursma *et al*., 2019). Two recent fully expanded leaves were sampled at pre-dawn from five replicate saplings per population × treatment. Leaves were scanned for leaf area and weighed immediately using a balance (0.0001 g) and measurement time was recorded. Leaves were then suspended in a growth chamber and allowed to slowly desiccate under stable conditions with temperature set at 20 °C, relative humidity at 75 %, light to 600-2000 µmol m^-2^ s^-1^ and fan speed to 200 m^2^s^-1^. Leaf mass and measurement time were recorded hourly on the first day and two measurements were taken on the second day, three hours apart. Leaves were then dried at 70 °C for at least 48 hours and weighed to obtain leaf dry mass.

Changes in leaf mass with time had an initial exponential decay relationship with high water loss prior to *P_gs90_*. Minimum leaf conductance (g_min_) was calculated from the slope of the latter linear region of the relationship between decreasing leaf mass (g) and increasing time (mins). This was converted from g^-1^ min^-1^ to mmol m^-2^ s^-1^ by dividing by projected leaf area (m^-2^) and the mean chamber VPD, (approx. 0.5 kPa), and converting the mass loss from g to mmol H_2_O.

**Figures**

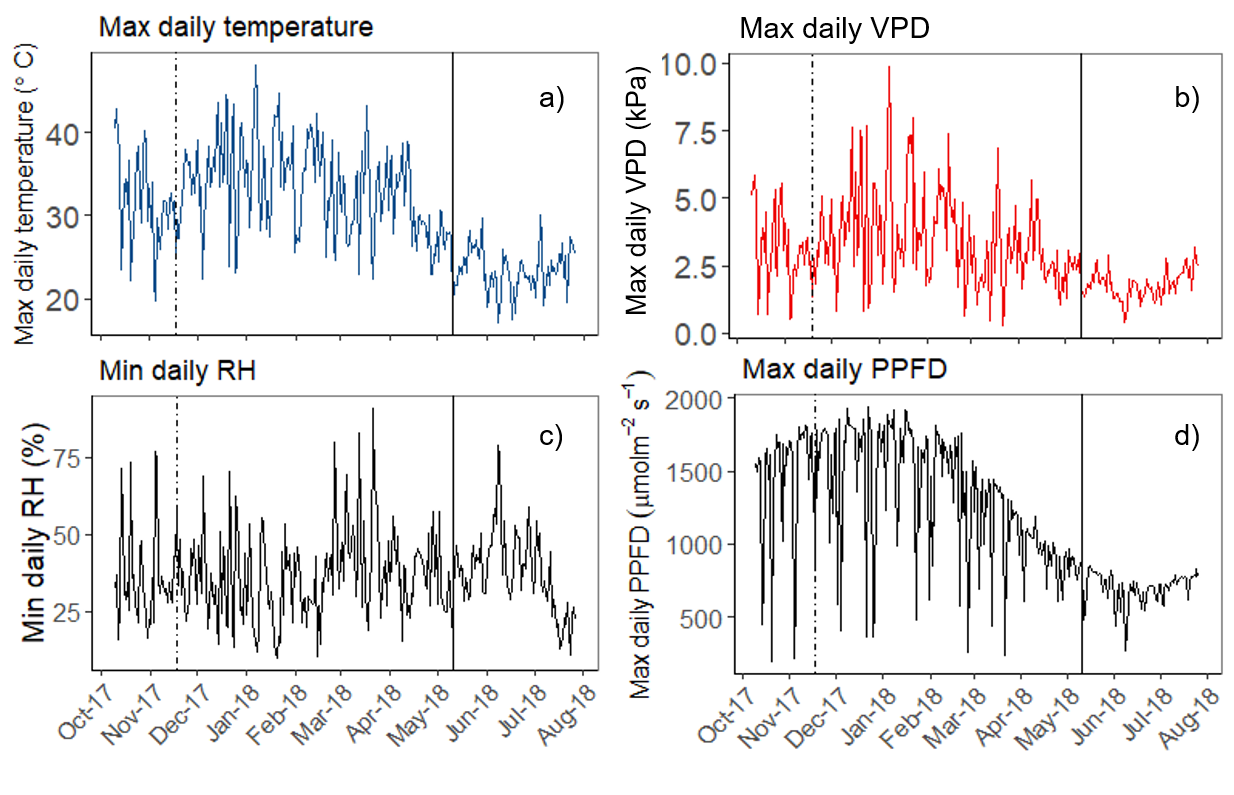

Figure S1. Maximum daily air temperature (°C, a), maximum daily vapour pressure deficit (VPD, kPa, b), minimum daily relative humidity (RH, %, c) and maximum daily photosynthetic photon flux density (PPFD, umolm-2s-1, d) in the poly-tunnel over the duration of the experiment. The broken vertical line indicates the date treatments commenced. The solid vertical line indicates commencement of the *drought to critical failure*

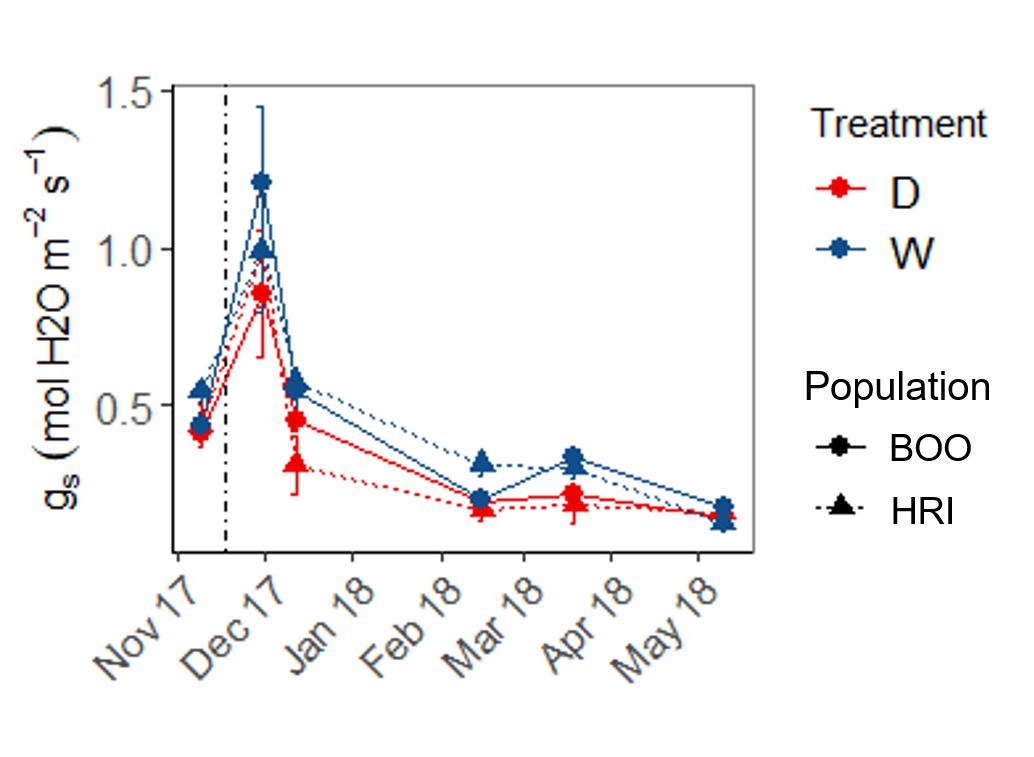

Figure S2. Mean stomatal conductance (*g_s_*, mol H_2_O m^-2^s^-1^) over time during the end of the *establishment phase* and the *treatment phase* of the experiment in Hill River (HRI, dashed line, triangle) and Boorara (BOO, solid line, circle) saplings grown under the well-watered (W, blue) and water deficit (D, red) treatments. Mean ±1SE are presented. The vertical dashed line indicates the date when treatments commenced.

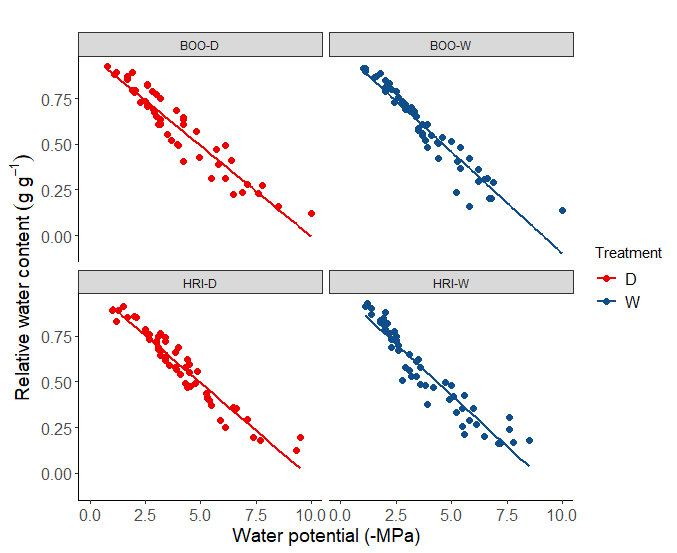
Figure S3. Relationships between relative water content (g g-1) and stem water potential (-MPa) of branches dried down in the laboratory in Boorara (BOO) and Hill River (HRI) populations grown under the well-watered (blue) and water deficit (red) treatments with linear models fitted for population × treatment combinations (r2 value BOO-D = 0.94, BOO-W = 0.94, HRI-D = 0.95, HRI-W = 0.93).

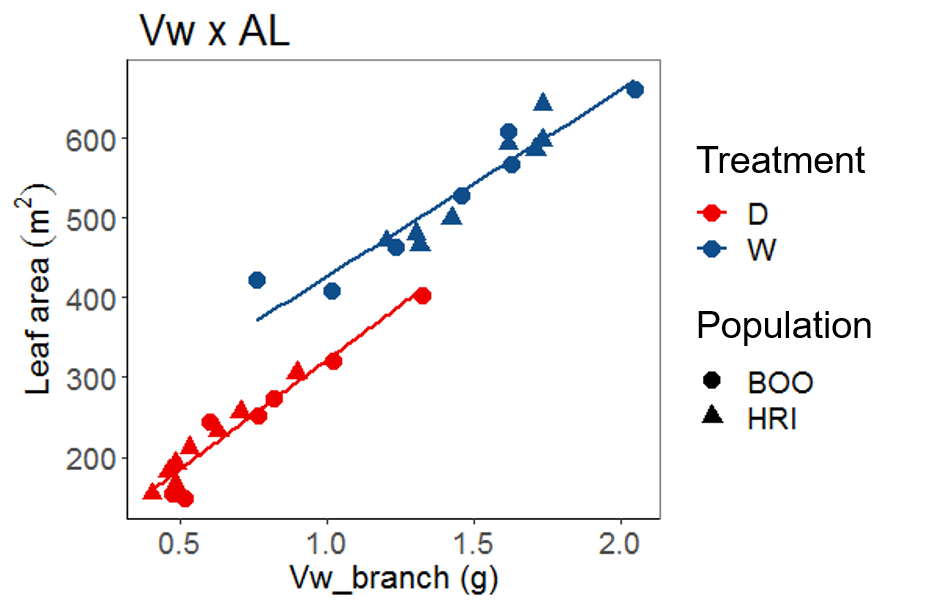

Figure S4. Relationship between total projected leaf area (*A*_L_) and total water storage (*V*_w_) across well-watered (W, blue) and water deficit (D, red) treatments including data from both Boorara (BOO, circles) and Hill River (HRI, triangles) populations. Lines indicate linear relationships for treatments with populations pooled (r^2^ D = 0.94, W = 0.90; *P* < 0.000).

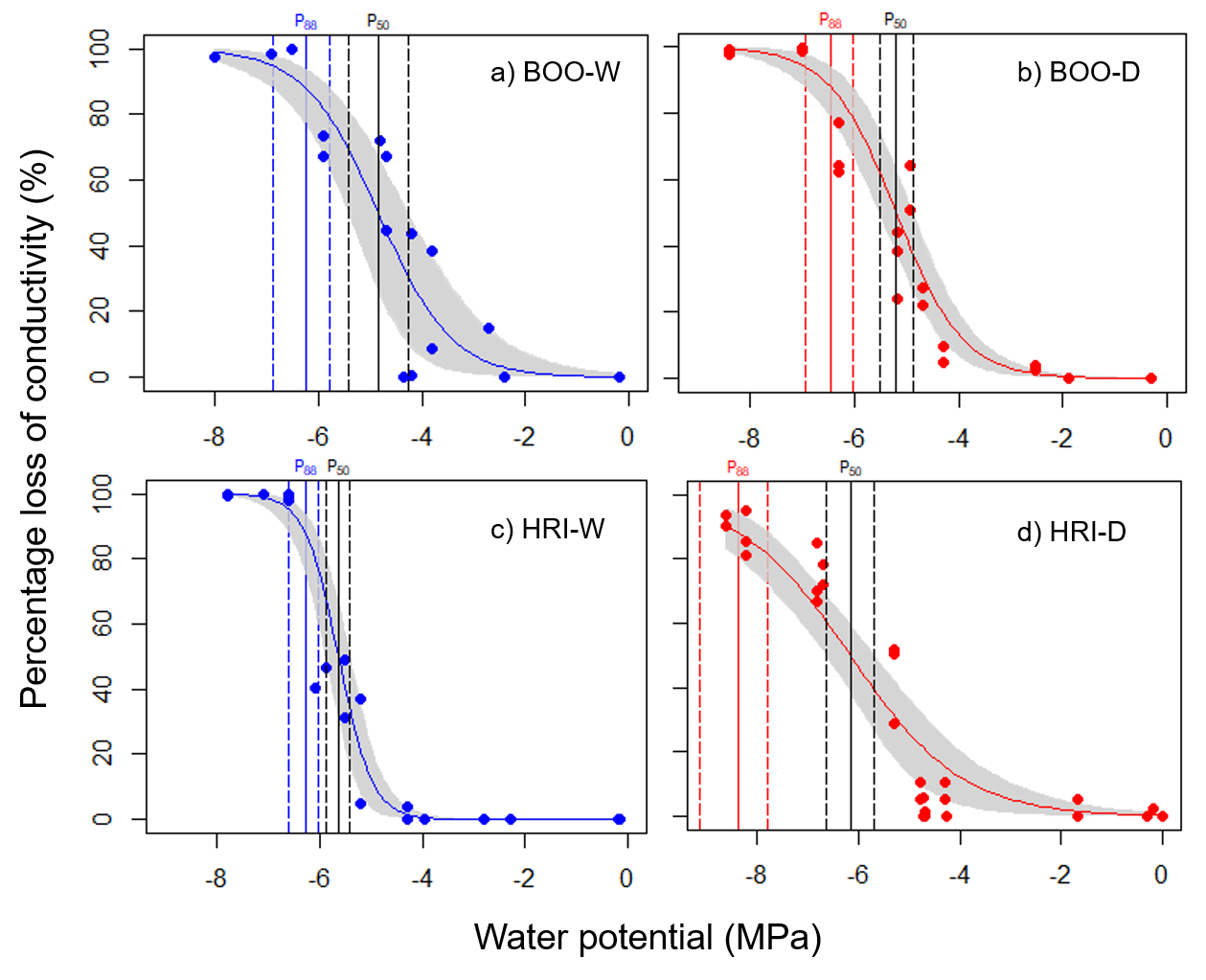

Figure S5. Percentage loss of conductivity curves for Boorara (BOO) and Hill River (HRI) populations grown under well-watered (W, blue) and water deficit (D, red) treatments. The water potentials at 50 % (*P*_50_, black lines) and 88 % (*P*_88_, coloured lines) loss of hydraulic conductivity are shown by vertical solid lines with 95 % confidence intervals indicated by the dashed lines

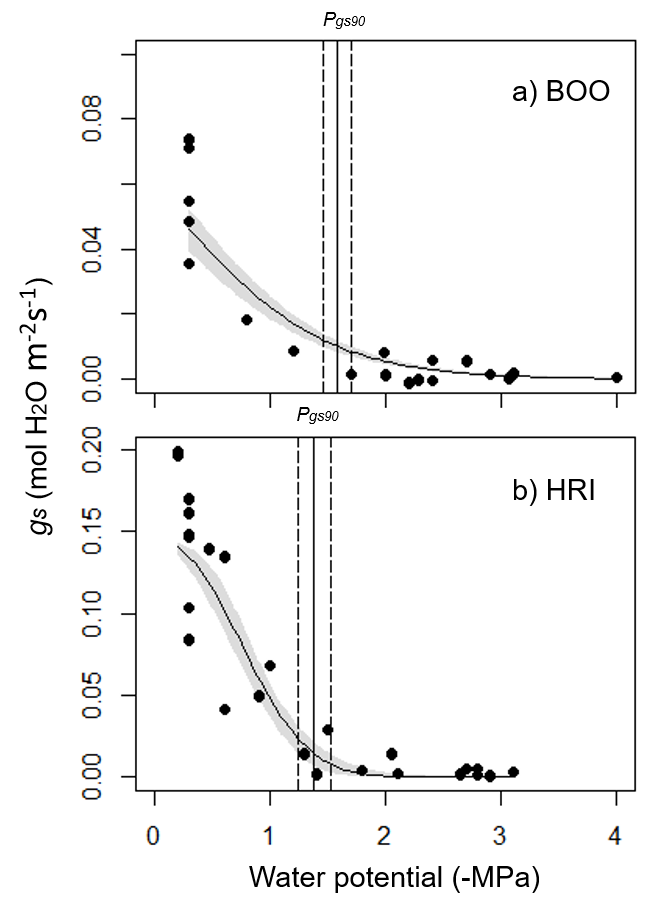

Figure S6. Stomatal closure curves showing the relationship between stomatal conductance (*g_s_*, mol H_2_0 m^-2^s^-1^) and leaf water potential (-MPa) in Boorara (BOO, a) and Hill River (HRI, b) saplings. The point of stomatal closure at 90 % loss of conductivity (*P_gs90_*) is indicated by the solid vertical line with 95 % confidence intervals shown by the vertical dashed lines.
